# Regular Tai Chi Practice Is Associated with Improved Memory as well as Structural and Functional Integrity of the Hippocampal Formation in the Elderly

**DOI:** 10.1101/2020.07.26.222190

**Authors:** Chunlin Yue, Qian Yu, Yanjie Zhang, Fabian Herold, Jian Mei, Zhaowei Kong, Stephane Perrey, Jiao Liu, Notger G. Müller, Zonghao Zhang, Yuliu Tao, Arthur Kramer, Benjamin Becker, Liye Zou

## Abstract

**Objective:** The current study aimed at determining effects of Tai Chi as an example of a combined motor-cognitive exercise relative to regular walking as an example of an exercise without cognitive demands on cognitive functioning and the functional and structural integrity of the brain in the elderly.

**Methods:** Healthy elderly women with at least 6 years of regular Tai Chi or brisk walking exercise were recruited and underwent cognitive assessment via the Montreal Cognitive Assessment and brain structural and resting state functional MRI assessments.

**Results:** Episodic memory in Tai Chi group was superior to that of the walking group; (2) higher gray matter density in inferior and medial temporal regions, including the hippocampal formation; (3) higher ReHo in temporal regions, specifically the fusiform gyrus and hippocampal formation (4) significant partial correlations were found between the gray matter density of the left hippocampus and episodic memory in the whole sample (5) significant partial correlations were observed between the ReHo in left hippocampus, left parahippocampal, left fusiform and delayed memory task was observed among all subjects.

**Conclusion:** The present study suggest that long-term Tai Chi practice may improve memory performance via remodeling structure and the function of the hippocampal formation.

## INTRODUCTION

During the last decades the number of older individuals in the society increases worldwide. In this population subjective and objective cognitive decline are highly prevalent and both have been associated with an increased risk for developing dementia, particularly in those over 65 years of age (Qiu and Fratiglioni, 2018). Age-related cognitive impairments are often pronounced in the domain of learning and memory, specifically episodic memory that binds personal experience with the spatio-temporal environment (Tromp et al., 2015). As one of the most important cognitive components, episodic memory is associated with recollection of personal experiences related to when and where an event takes place (Tulving, 1983), which is necessary for both work performance and daily life activity. Episodic memory represents a critical building block for successful everyday functioning and critically relies on the integrity of medial temporal lobe structures, specifically the hippocampal formation (Becker et al., 2013; Davachi and Wagner, 2002). Moreover, episodic memory is particularly sensitive to brain aging (Nyberg et al., 2012; Shing et al., 2010), and usually the first memory system to decline in both normal and pathological aging (Tromp et al., 2015). Additionally, age-dependent episodic memory decline has been observed to be accompanied by volume reduction in medial temporal and to a lesser extend frontal brain regions (Persson et al., 2006; Rajah et al., 2010; Van Petten et al., 2004).

Given the critical roles of memory dysfunction in everyday functioning, including not only cognitive but also social domains (Allen and Fortin, 2013), timely interventions (i.e., physical activity) are needed to postpone or counteract age-related memory decline. Accumulating evidence from different lines of research suggests that physical activity triggers a multitude of neuroprotective and neurorestorative effects on the brain which in turn promote cognitive performance (Hillman et al., 2008; Yu et al., 2020). For instance, walking - which represents the most common and favorite activity among the elderly (Szanton et al., 2015) - has been shown to be associated with better general cognitive functioning, specifically memory and decreased likelihood to develop dementia (Yaffe et al., 2001).

Tai Chi, a multicomponent mind-body exercise, combines slow physical activity with relaxation to serve as a movement meditation (Wang et al., 2016). Prior trials suggested that the beneficial effects of Tai Chi are mediated by a physical component which capitalizes on the benefits of physical exercise and a mind component which additionally promotes psychological well-being, life satisfaction and improved perception of health (Black et al., 2014; Wang et al., 2016). With respect to cognitive performance, numerous studies have demonstrated the cognitive benefits (including memory) of Tai Chi on the behavioral level (Ji et al., 2017; Lim et al., 2019; Sungkarat et al., 2018; Zou et al., 2019), however, the potential neurobiological processes and relevant brain mechanisms (structure and function) are still not-well understood (Chen et al., 2020; Cui et al., 2019; Wei et al., 2013; YueZhang et al., 2020; YueZou et al., 2020). In addition, it is still unclear whether Tai Chi differentiates from purely physical exercise (e.g. walking) with respect to the beneficial effects on cognition and the underlying neurobiological mechanism. In order to better understand cognitive benefits of Tai Chi in the elderly, we aimed at identifying cognition, brain functional and structural profiles differences between regular Tai Chi practitioners and well-matched elderly walkers.

## METHODS

### Study Participants

Participants were recruited in the Suzhou community and the Suzhou Sports Bureau via word-of-mouth advertisements and posters. A total of 200 older individuals were screened according to the following study criteria to identify eligible participants: (i) right-handed as measured by the *Edinburgh* Handedness Questionnaire (Oldfield, 1971), (ii) normal visual acuity or corrected vision, (iii) age 60 years or above, (iv) successful completion of the primary school, (v) normal cognitive performance indicated by a score of 26 or higher in the Montreal Cognitive Assessment Scale (MOCA), (vi) more than 6 years of Yang-style Tai Chi training with, at least, 5x 90-minute sessions per week and an award in Yang-style Instruction Certificate (Tai Chi group) or walking exercise for more than 6 years, at least, 5 times weekly while each session lasted, at least, 90 minutes (Walking group) (YueZhang et al., 2020; YueZou et al., 2020). Volunteers were excluded if they had: (1) diagnosis of a mental and/or physical illness; (2) history of neurodegenerative diseases or brain disorder; (3) drug and alcohol addiction; (4) contraindications for MRI. 46 participants were included for magnetic resonance imaging conducted in the Brain Research Center of the Second Affiliated Hospital of Soochow University. All participants provided written informed consent and all study procedures were in accordance with the latest revision of the Declaration of Helsinki. Four individuals were excluded from data analysis due to excessive head movement (n = 3) and claustrophobia (n = 1), leading to a final sample size of 20 and 22 in the Tai Chi group (TCG) and walking group (WG), respectively.

### Assessment of Cognitive Functioning

The MOCA, administrated by trained research assistants, was used to determine group differences on global cognition and its subdomains. This assessment tool is a paper and pencil instrument that takes about 10 minutes to administer and consists of the following measurements: orientation, short-term memory, long-term memory, visuospatial skills, attention, language, verbal fluency, calculation and abstraction. The MOCA has excellent internal consistency (r = 0.83) and has been validated in 56 languages to assess cognitive performance globally (Nasreddine et al., 2005).

### Data Acquisition

A 3Tesla whole body magnetic resonance imaging (MRI) system (Philips Ingenia) was employed to collect the data. For each participant, we conducted one functional MRI resting-state assessment using gradient echo planar imagine (EPI) sequence with the following scan parameters (FOV: 220mm×220mm, TR: 2000ms, TE: 30ms, slices: 36, flip: =90°, slice thickness: 4mm, matrix size: 64×64, total scan time: 400s. For the brain structural assessments each participant additionally underwent acquisition of T1-weighted images using a magnetization-prepared rapid gradient echo sequence (MPRAGE) with the scan parameters: voxel size: 0.625×0.625×1mm, TR: 7.1ms, TE: 2.98ms, flip: 9°, slice thickness : 1.0mm, FOV: 256×256mm, matrix size: 256×256, scanning layer by layer). MRI acquisition was conducted by an experienced radiologist. Each participant was asked to lie comfortable and to think of nothing in particular while moving their head and body as little as possible. Participants were explicitly requested to stay awake during data acquisition.

### Data Processing

#### Voxel-Based Morphometry (VBM)

Structural MRI data was analyzed and processed in Matlab 2013b (The Mathworks®, Natick, MA, USA). A SPM12 (Statistical Pararmetric Mapping; https://www.fil.ion.ucl.ac.uk/spm/) and CAT12 (Computational Anatomy Toolbox; http://www.neuro.uni-jena.de/cat12/CAT12). The recommended default parameters were employed to preprocess the structural image data. The 3D structural image of each participant were initially segmented into gray matter (GM), white matter (WM), and cerebrospinal fluid (CSF). Voxel resolution was set to 2*2*2mm. Spatial smoothing was performed using a Gaussian kernel with a full width at half maxima (FWHM) of 8 mm. To avoid a partial density effect in the boundary between GM and WM all voxels with GM value below 0.2 were excluded. Total intracranial volume (TIV) was calculated and considered as a covariable in all further statistical analysis.

#### Regional Homogeneity (ReHo) Analysis

The RESTplus software was used to preprocess the resting-state fMRI data in Matlab2013b (Jia et al., 2019).Data preprocessing included standard processing steps, including: (1) deletion of the first 1-time points, and the data of 190 time points in the resting-state were retained; (2) time layer and head motion correction according to the realign curve. Participants with head motion at x, y and z-axis translation greater than 3 mm and rotation greater than 3° were excluded; (3) registration of the motion-corrected to the Montreal Neurological Institute (MNI) space via normalization of the functional time-series MNI EPI template to account for interindividual anatomical differences; (4) Finally, a linear regression was used to remove linear drift, white matter signals, CSF signals, and 24 head parameters were regressed as covariables to remove physiological influences (Murphy et al., 2009); (5) temporal band-pass filtering (0.01-0.08 Hz). Based on Kendall’s coefficient of Concordance (KCC), the ReHo value of each voxel was subsequently calculated using RESTplus software to obtain whole brain ReHo maps of each subject was obtained. For standardization purpose, each ReHo map was divided by the average ReHo value of the entire brain. A Gaussian kernel of 8 mm FWHM was used for spatial smoothing (Zang et al., 2004).

#### Statistical Analyses

The SPSS 19.0 software (SPSS Inc., Chicago, IL, USA) was used for data analysis. A T test was performed to compare differences between measures obtained from the Tai Chi and walking groups, and the level of significance was set at 0.05. Cohen’s d (effect size) was classified as small (d = 0.2), medium (d = 0.5), and large (d ≥ 0.8). The independent-samples T test in SPM software was used to analyze the differences on GM and ReHo between the Tai Chi and walking groups. Analysis with a threshold of voxel-level *p* < 0.001 and cluster-level *p* < 0.01 GRF (Gauss Random Field) correction was applied for the ReHo and VBM analyses. A partial correlation analysis was used to compare the correlation between sub-items of MOCA and GM and ReHo, with age (years), sex, and education (years) as covariates. We used the Bonferroni correction to control for multiple comparisons. The correlation coefficients were defined as follows: 0.00 to 0.19 no correlation; 0.20 to 0.39 low correlation; 0.40 to 0.59 moderate correlation; 0.60 to 0.79 moderately high correlation; 0.8 high correlation (Zhu, 2012). The Xjview (http://www.nitrc.org/projects/bnv) (Xia et al., 2013) and Origin 9 software were used to present the results.

## RESULTS

Independent sample T tests revealed no statistically significant differences between the two groups for age, years of education or global cognitive performance (operationalized by MOCA scores) (details see **Table 1**). Examining the cognitive sub-domains assessed by the MOCA revealed a significant and specific between-group difference in the domain of delayed recall with the TCG performing significantly better than WG (details see **Table 2**). The groups did not differ in any cognitive domain, suggesting highly specific differences in the domain of episodic memory.

**TABLE 1.** Characteristics of the participants in TCG and WG

|  | TCG (N=20) | WG (N=22) | *T* | *p* |
| --- | --- | --- | --- | --- |
| Age (year) | 62.9±2.4 | 63.27±3.6 | -0.393 | 0.193<sup>a</sup> |
| Education (year) | 9.05±1.8 | 8.86±2.74 | 0.188 | 0.074<sup>b</sup> |
| Handedness (left/right) | 0/20 | 0/22 | - | - |
| Year of practice | 16.58±7.33 | 14.95±5.94 | 0.823 | 0.42<sup>a</sup> |
| Amount of exercise ( h/week ) | 9.91±3.04 | 10.87±2.06 | -1.26 | 0.20<sup>b</sup> |
| MOCA | 28.4±1.5 | 27.5±1.5 | 1.94 | 0.83<sup>b</sup> |
| CSF ( cm<sup>3</sup> ) | 356.30±142.11 | 327.36±103.56 | 0.74 | 0.46<sup>b</sup> |
| GMV ( cm<sup>3</sup> ) | 560.45±61.73 | 533.91±45.08 | 1.56 | 0.13<sup>b</sup> |
| WM ( cm<sup>3</sup> ) | 483.50±64.27 | 462.27±41.53 | 1.25 | 0.22<sup>b</sup> |
| TIV ( cm<sup>3</sup> ) | 1400.35±15.10 | 1323.77±142.53 | 1.43 | 0.16<sup>b</sup> |
| GM/TIV | 0.40±0.04 | 0.41±0.03 | -0.16 | 0.87<sup>b</sup> |
| GM+WM ( cm<sup>3</sup> ) | 1043.95±119.27 | 996.18±78.81 | 1.51 | 0.14<sup>b</sup> |
| GM/GM+WM | 0.54±0.02 | 0.54±0.02 | 0.25 | 0.81<sup>b</sup> |

*Note: a. Two sample T-test (two-tailed); b. Nonparametric test (Mann–Whitney test); MOCA: Montreal Cognitive Assessment; GMV: Gray matter volume; CSF: Cerebrospinal fluid; WM: White matter; TIV: Total intracranial volume.*

**TABLE 2.** Comparisons of cognitive domains of the MOCA between TCG and WG

|  | TCG<br>(n = 20 ) | WG<br>(n = 22 ) | *p* | Cohen's *d* |
| --- | --- | --- | --- | --- |
| MOCA | 28.4±1.5 | 27.5±1.5 | 0.83<sup>a</sup> | 0.600 |
| Visuospatial | 1.6±0.60 | 1.82±0.39 | 0.167<sup>a</sup> | -0.435 |
| Draw the clock | 2.7±0.47 | 2.86±0.35 | 0.213<sup>a</sup> | -0.386 |
| Naming | 3±0 | 2.64±0.73 | 0.029<sup>a</sup> | 0.674 |
| Attention | 5.4±0.68 | 5.5±0.67 | 0.635<sup>a</sup> | -0.148 |
| Repeat sentence | 1.7±0.57 | 1.82±0.39 | 0.445<sup>a</sup> | -0.246 |
| Verbal fluency | 1±0 | 1±0 |  |  |
| Abstract ability | 1.9±0.31 | 1.82±0.39 | 0.456<sup>a</sup> | 0.227 |
| Delayed recall | 4.45±0.89 | 3.32±1.25 | 0.002<sup>a*</sup> | 1.041 |
| Orientation | 5.85±0.37 | 5.95±0.21 | 0.273<sup>a</sup> | -0.332 |

*Note: a. Nonparametric test (Mann–Whitney test); \* indicates significant differences (*p*<0.01).*

### VBM Analysis

The VBM analysis of the T1-weighted structural images showed that gray matter density in the left cerebellum and right inferior and left medial temporal lobe regions, including the left hippocampus and parahippocampal gyrus was significantly higher in the TCG group as compared to the WG group (Table 3, Figure 1). Then performed a whole-brain voxel-wise between-subject ANOVA on the VBM maps in two Groups (TCG, WG) to detect regions showing intervention-related difference. The threshold for significant difference was set to p<0.01 cluster mass-level GFR corrected with a cluster building threshold of uncorrected p = 0.001 on voxel level. In further studies, the regions showing some significant differences were defined as regions of interest (ROIs). We extracted the mean gray matter density signal in each ROI and used partial correlation analysis to examine the correlation between gray matter density signal and delayed memory score of the two groups.

**TABLE 3.** Brain regions with higher gray matter density in TCG as compared to WG

| Brain Region | MNI coordinates, mm |  |  | Voxels, n | *Z*<sub>peak</sub> | *p* (voxel) | *p* (cluster) |
| --- | --- | --- | --- | --- | --- | --- | --- |
|  | x | y | z |  |  |  |  |
| Cerebelum_8_L | -34.5 | -61.5 | -49.5 | 310 | 4.26 | 0.001 | 0.01 |
| ITG_R | 51.5 | -5.5 | -27.5 | 204 | 4.05 | 0.001 | 0.01 |
| HIP_L/ PHG_L | -26.5 | -35.5 | -7.5 | 197 | 4.13 | 0.001 | 0.01 |

Note: MNI coordinate values were calculated based on the standard template space of the Montreal Institute. *Cerebelum\_8\_L*: left inferior cerebellum; *ITG\_R*: right inferior temporal gyrus; *HIP\_L*: left hippocampus; *PHG\_L*: left parahippocampal gyrus.

**FIGURE 1.**
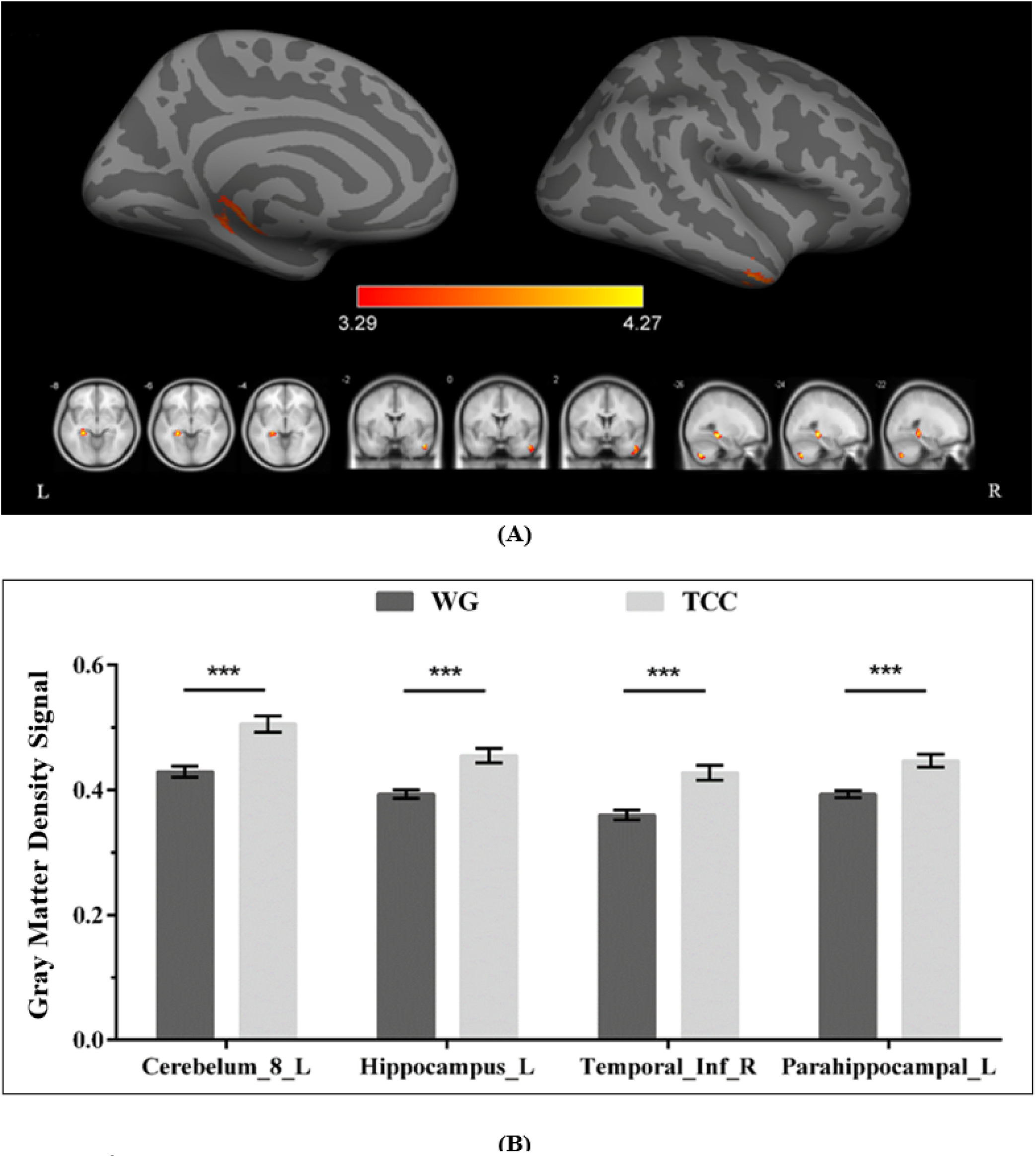
Regional gray matter differences as determined by VBM between TCG and WG group. (A)The threshold for significant changes was set to *p* < 0.01 cluster mass-level GFR corrected with a cluster building threshold of *p* = 0.001 uncorrected on voxel level. The warm color indicates that the GMV of the TCG was greater than that of WG. (B)Bar plots showed the mean gray matter density signal in these ROIs for TCG and WG group. The independent sample t-test was used to compare the effects of two groups. *** means *p*<0.001. *Abbreviations. Cerebelum_8_L: left inferior cerebellum; ITG_R: right inferior temporal gyrus; HIP_L: left hippocampus; PHG_L: left parahippocampal gyrus*.

### Reho Analysis

Resting-state fMRI analysis revealed significantly higher regional ReHo activation in the TCG group in the left medial temporal lobe, specifically the hippocampus and parahippocampus, as well as the fusiform gyrus relative to the WG group (**Table 4, Figures 2**). Then performed a whole-brain voxel-wise between-subject ANOVA on the ReHo maps in two Groups (TCG, WG) to detect regions showing intervention-related difference. The threshold for significant difference was set to p<0.01 cluster mass-level GFR corrected with a cluster building threshold of uncorrected *p* = 0.001 on voxel level. In further studies, the regions showing some significant differences were defined as regions of interest (ROIs). We extracted the mean ReHo value in each ROI and used partial correlation analysis to examine the correlation between ReHo and delayed memory score of the two groups.

**TABLE 4.** Brain regions with different ReHo in TCG and WG

| Brain regions | MNI coordinates,mm |  |  | Voxel,n | Z<sub>peak</sub> | p ( voxel ) | p ( cluster ) |
| --- | --- | --- | --- | --- | --- | --- | --- |
|  | x | y | z |  |  |  |  |
| HIP_L/ PHG_L/ FG_L | -27 | -33 | -3 | 280 | 5.34 | 0.001 | 0.01 |

*Note: MNI coordinate values were calculated based on the standard template space of the Montreal Institute. Abbreviations. HIP\_L: left hippocampus; PHG\_L: left parahippocampal gyrus; FG\_L: left fusiform.*

**FIGURE 2.**
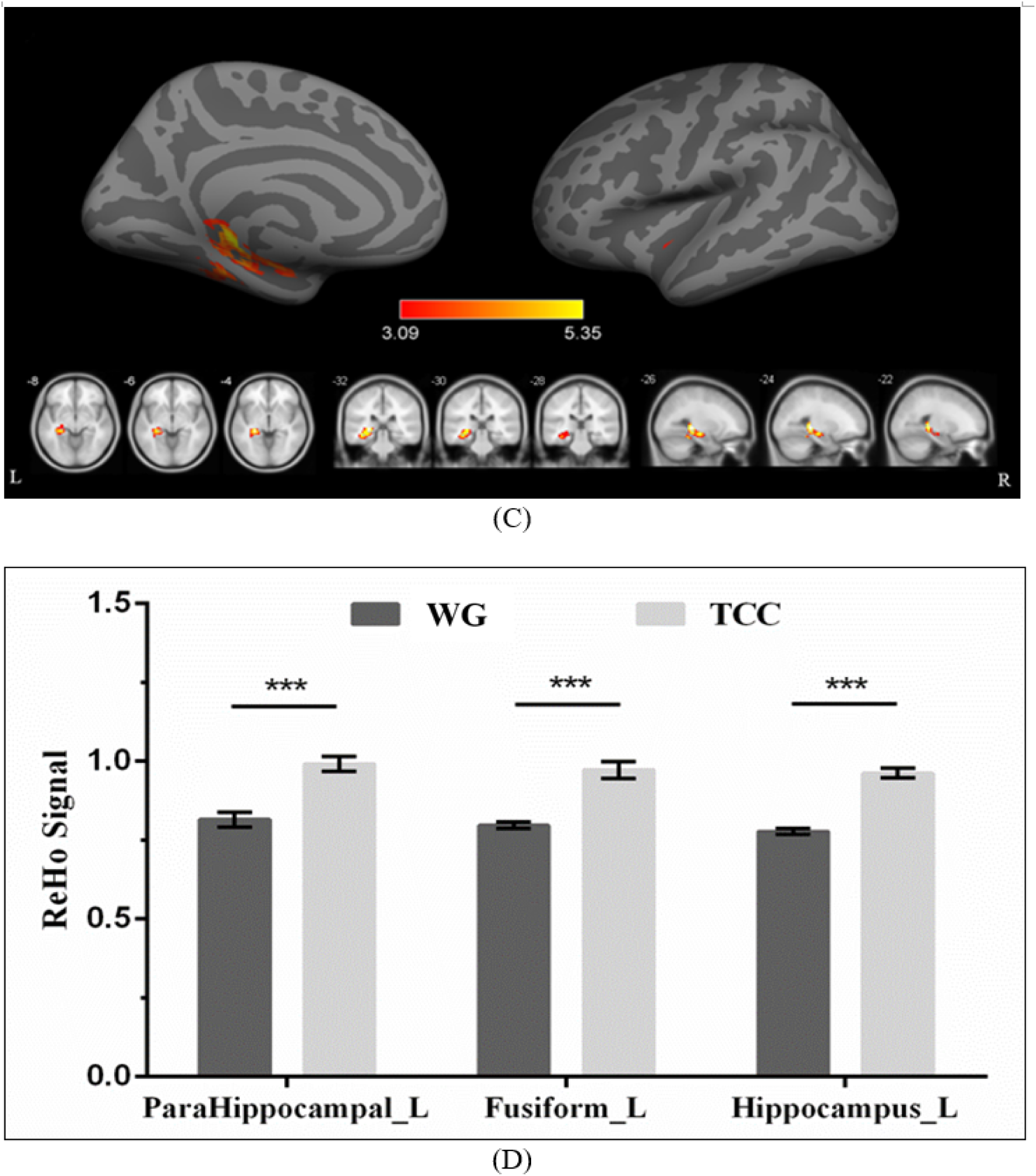
Regional differences in ReHo between TCG and WG. (C) ReHo analysis showed that the ReHo activations of left hippocampus, parahippocampal gyrus and fusiform in the participants who practiced Tai Chi were higher than those in WG. The threshold for significant changes was set to *p*<0.01 cluster mass-level GFR corrected with a cluster building threshold of *p* = 0.001 uncorrected on voxel level. (D) Bar plots showed the mean ReHo signal in these ROIs for TCG and WG group. The independent sample t-test was used to compare the effects of two groups. *\*\*\* means p<0*.*001. Abbreviations. HIP_L: left hippocampus; PHG_L: left parahippocampal gyrus; FG_L: left fusiform*.

### Correlation between Brain Structure, Brain Function and Behavioral Performance

Partial correlation analyses (controlling for the participants’ age and level of education) was conducted on the four regions of interest (ROI) (left inferior cerebellum, right inferior temporal gyrus, left hippocampus and parahippocampal gyrus) obtained by VBM analysis and nine domains of MOCA (Visuospatial, Draw the clock, Naming, Attention, Repeat sentence, Abstract ability, Delayed recall, and Orientation). As shown in **Figure 3**, we observed a significant moderate and positive correlation between the gray matter density in left hippocampus specifically for the delayed memory task (*r* =0.547, *p* < 0.01, Bonferroni correction) at a Bonferroni correction threshold of 0.0014 (0.05/36) when the whole sample was considered (Ludbrook, 1998). In the WG, the correlation between gray matter density in left hippocampus and delayed memory task failed to reach statistical significance at the Bonferroni corrected level (see Statistical analysis) and thus could be considered as only marginally significant (*r*=0.613, *p*s<0.1, Bonferroni correction; see Figure 3). On the other hand, no significant correlation between the gray matter density changes in the left hippocampus and delayed memory task was found in the TCG (*r*=0.241, *p*=0.307; see Figure **3**) (Ludbrook, 1998). Additionally, there was no significant correlation between the gray matter density of the remaining 2 ROI and other sub-items of the MOCA.

**FIGURE 3.**
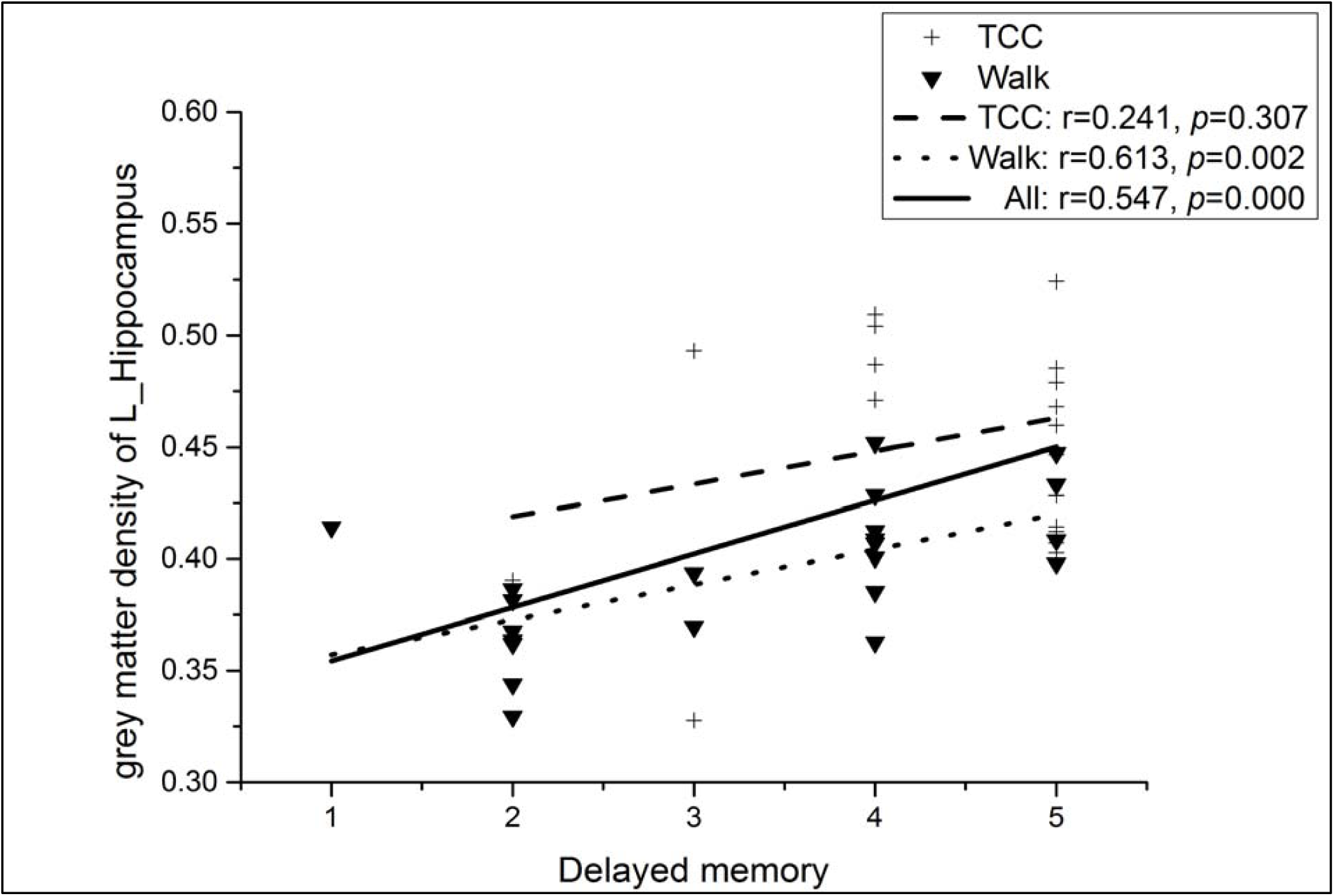
Correlation between gray matter density of left hippocampal and delayed memory

Partial correlation analysis was conducted on the three region of interest (ROI) (the left hippocampus, parahippocampal gyrus and fusiform) obtained by ReHo analysis and nine sub-items of MOCA (Visuospatial, Draw the clock, Naming, Attention, repeat sentence, Abstract ability, Delayed recall, and Orientation) with controlling the participants’ age and education (Ludbrook, 1998). A moderately high and positive correlation between the ReHo in left hippocampus(*r*=0.679, *p*<0.01, Bonferroni correction), left parahippocampal (*r*=0.612, *p*<0.01, Bonferroni correction), left fusiform (*r*=0.570, *p*<0.01, Bonferroni correction) and delayed memory task was observed when the whole sample was considered at a Bonferroni correction threshold of 0.0019 (0.05/27). In the WG, a significant moderately high and marginally significant correlation between the ReHo in left hippocampus and delayed memory task was observed (*r* =0.619, *ps*<0.1, Bonferroni correction; see **Figure 4**). On the other hand, no significant correlation between the ReHo changes in the left hippocampus(*r*=0.507, *p*=0.022; see Figure 4), left parahippocampal(*r*=0.448, *p*=0.048; see Figure 4), left fusiform(*r*=0.450, *p*=0.047; see Figure 4) and delayed memory task was found in the TCG (Ludbrook, 1998).

**FIGURE 4.**
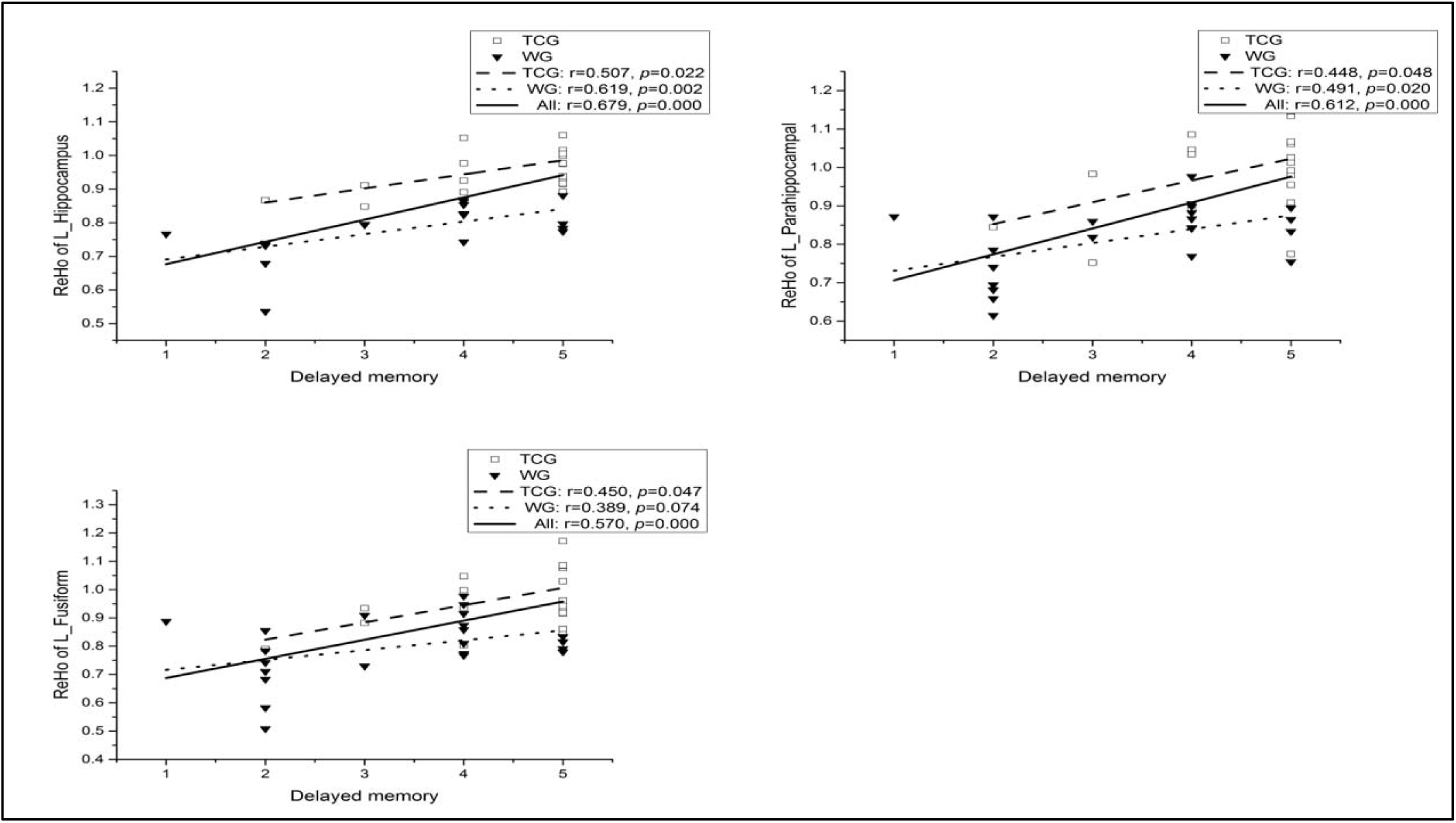
Correlation between ReHo of left hippocampal and delayed memory

## DISCUSSION

The current study employed a cross-sectional design to determine differences in cognitive performance and brain integrity between long-term elderly practitioners of Tai Chi and walking exercises. Compared with the walking group, Tai Chi practitioners performed better in the domain of delayed memory and additionally exhibited higher gray matter density in temporal regions, including the left hippocampus and the adjacent left parahippocampal gyrus. On the functional level the Tai Chi practioners exhibited higher spontaneous ReHo activation in similar temporal regions, including left hippocampus and parahippocampal gyrus as well as the fusiform gyrus as compared to the walking group. Finally, significant associations were observed between the left hippocampal gray matter density and delayed memory performance in the whole sample level, confirming an important contribution of this region to memory performance in the present sample. In the following sections, we will discuss these findings in more detail.

### Hippocampus: Structural Changes, Functional Changes and Relationship with Memory

Long-term Tai Chi practitioners showed higher density and ReHo activation in the hippocampus and parahippocampal gyrus as compared to the WG. Moreover, on a behavioral level, we observed that long-term Tai Chi practice was more effective than walking exercise in enhancing selected aspects of cognitive function (i.e., long-term memory) which is in accordance with previous observations (Ji et al., 2017). This observation fits well with the observations of a positive association between better memory performance and higher density of the hippocampus (Erickson et al., 2011). The episodic memory performance reflects the ability to recollect past experience in the temporal and spatial context, which is highly influenced by structure of the temporal lobe (especially in hippocampus) (Grilli et al., 2018). More specifically, this brain area is critically engaged in consolidation and retrieval, particularly associated information. Aging-related hippocampal shrinkage (atrophy) has been increasingly reported and has been associated with memory impairments and a strongly increased risk for the development of dementia and Alzheimer’ disease (Jack et al., 2010; Small et al., 2011). Previous studies consistently indicated that hippocampal neuroplasticity (neurogenesis) could be induced by regular physical exercise, especially physical training. Such exercise-induced biochemical changes in the brain region have been observed to associate with (1) increased serum level of brain-derived neurotrophic factor (BDNF) that is conducive to long-term potentiation and proliferation of neurons; (2) vascular endothelial growth factor (VEGF) that is beneficial for blood vessel survival and growth; (3) insulin-like growth factor (IGF)-1 that can contribute to several neural and angiogenic process (Cotman et al., 2007). For example, a randomized controlled trial by Erickson et al (Erickson et al., 2011) indicated that 1-year aerobic exercise training effectively contributed to 2% increase of hippocampal volume that corresponded to improved spatial memory performance and such increased size in this brain region was positively associated with serum levels of BDNF, whereas older adults in the control group demonstrated nearly 1.5% hippocampal volume loss within this intervention period.

Additionally, a seminal study found that a 10-min bout of mild exercise effectively improved pattern separation and functional connectivity between hippocampal dentate gyrus/CA2 and surrounding brain regions and such objectively measured results were positively associated with memory performance (Suwabe et al., 2018). Although the above-summarized evidence suggest that the observed greater hippocampal density could be triggered by long-term Tai-Chi practice, our cross-sectional study design allows limits direct conclusions with respect to causal relationships and thus further longitudinal study are required to confirm this assumption empirically. Furthermore, it is widely accepted that ReHo of the resting-state signals play a critical role in cognitive function (including learning and memory) (Zang et al., 2004). Age-associated reduction of ReHo in several brain regions was negatively associated with behavioral outcomes (Diciotti et al., 2017). Recently, a combined program that consisted of cognitive training, Tai Chi, and group counseling leads to a reorganization of ReHo patterns in several brain regions (including superior and middle temporal gyri), which was coincident with improved cognitive function after the 6-week intervention period (Zheng et al., 2015). Given that we observed greater ReHo of the left hippocampal area in the Tai-Chi group and that we noticed a positive neurobehavioral relationship between ReHo patterns and memory performance, it seems to be reasonable to assume that engaging in long-term Tai Chi exercise (in comparison to walking), is a valuable strategy to prevent the decline in cognitive performance as it preserves underlying neural correlates.

### Other Related Brain Regions in This Study

As compared to the WG, greater gray matter density in TCG was also observed in other brain regions, including the parahippocampal gyrus, cerebellum, and inferior temporal gyrus; higher ReHo activations were also found in parahippocampal gyrus and fusiform.

The parahippocampal gyrus, located just inferior to the hippocampus, plays an important and distinctive role in the processing of memory (Nemanic et al., 2004). The parahippocampal gyrus is comprised of the entorhinal, perirhinal and parahippocampal cortices (Burwell, 2000): cognitive information is usually collected by the perirhinal and parahippocampal cortices before being further processed in entorhinal cortex and hippocampus (Loprinzi, 2019). While the entorhinal cortex and entorhinal cortex process spatial and object-recognition information (Raslau et al., 2015), the parahippocampal cortex has a critical role in episodic memory (Aminoff et al., 2013). Moreover, in the previous studies (Loprinzi, 2019; Nemanic et al., 2004; Siddarth et al., 2018), extensive memory-related effects (i.e., increased neural excitability, blood flow, regional glucose metabolism, gray/white matter and functional connectivity) induced by physical exercise have been reported in the parahippocampal gyrus, which implies that exercise interventions like Tai Chi may facilitate the memory function through regulation of parahippocampal gyrus at molecular, structural and functional levels.

The major function of cerebellum was believed to be efficiently coordinating the timing and forces of multiple muscle groups to initiate voluntary movements, according to both anatomic and clinical evidence before the 1990s. However, later findings have cast doubt on this view. Cerebellar activation has been increasingly investigated using imaging techniques and found to associate with cognitive and affective tasks. Furthermore, among the patients whose injury limits to the cerebellum, a variety of non-motor symptoms have been recognized, especially those with cognitive affective syndrome (Doya, 2000; Schmahmann and Sherman, 1998). Evidences from functional mapping of the cerebellum showed that more than half of the cerebellar cortex is connected to the cognition-related areas of the cerebral cortex (Buckner et al., 2011). It is worth noting that the anterior cerebellum (lobules I-IV and V) and anterior part of lobule VI are mainly related with motor functions (Grodd et al., 2001; Nitschke et al., 1996; Rijntjes et al., 1999) while the posterior part plays an evident role in neurocognition and emotion (Eekers et al., 2018). Thus, cerebellum may contribute to physical and cognitive effects induced by Tai Chi through different regions and connections.

Regarding the fusiform gyrus, a previous study using MRI indicated that fusiform gyrus area as part of human brain is responsible for face-specific processing. Furthermore, greater activation in this brain region was observed during facial stimuli as compared to objects (McCarthy et al., 1997). Haxby and colleague proposed a model for face perception and emphasized the distributed human neural system. Specifically, the face-responsive region in the fusiform gyrus was observed to explain more for the unchanging features of a specified facial identity, whereas the face-responsive region in the superior temporal sulcus could be involved in changeable aspects (name) of faces (Haxby et al., 2000). Thus, the fusiform gyrus plays a critical role in social communication or interaction that is associated with cognitive benefits. This assumption is supported by an experimental study suggesting that community-based Tai Chi training facilitates social interaction, which, in turn, might contribute to the observed increase in brain volume and behavioral performance in non-demented older adults (Mortimer et al., 2012).

Regarding temporal gyrus, the primary function is connected with visual stimuli processing (Lafer-Sousa and Conway, 2013), which is involved with memory, perception and recognition (Bryan Kolb, 2014). Previous studies believed that temporal gyrus works in the memory processing through cooperation with other brain regions like hippocampus (key for storing the memory), parahippocampal gyrus (participation in differentiating between scenes and objects) and fusiform gyrus (dealing with facial and body recognition) (Gross, 2008; Spiridon et al., 2006). Additionally, in the study of Makizako et al. (Makizako et al., 2013), evidence suggested that higher exercise capacity is associated with better performance in logical and visual memory; and poor memory function was correlated with decreased gray matter volume in temporal gyrus, hippocampus and occipital gyrus. These outcomes indicated that exercise (like Tai Chi) may improve visual-stimuli-related memory through maintenance of gray matter volume of hippocampus (main region for memory), temporal gyrus and other brain regions, which is consistent with findings in our study.

### Strengths and Limitations

The strengths of this investigation are the high practical relevance of our findings as we used state-of-art neuroimaging to investigate the effects of popular physical exercises (Tai-Chi and walking) in a cohort vulnerable to cognitive decline. Furthermore, we have applied rigorous inclusion criteria allowing us to identify a cohort of long-term practitioners of Tai-Chi and walking which have, at least 10 years of experience. One limitation of this investigation, however, is that we did not account for some other potentially important factors (e.g., cardiorespiratory fitness) that might bias our findings. For instance, there is evidence in the literature suggesting that cardiorespiratory fitness level is associated with hippocampal volume (Erickson et al., 2009; Szabo et al., 2011) although this finding is not universal (Niemann et al., 2014). Additionally, as only elderly women were included in this study, it is unknowable whether our outcomes are applicable to people with different genders.

## CONCLUSION

In conclusion, in this cross-sectional study, we demonstrated that relative to walking Tai Chi was more effective in enhancing memory performance in a sample of healthy older Chinese women. The observed changes in hippocampal structure and function and significant behavioral relationships between hippocampal integrity and memory functions suggest that physical activity like Tai Chi is likely to benefit memory function via remodeling of hippocampal structure and function. Furthermore, our findings support the conclusion that engaging in Tai-Chi might be superior in delaying cognitive decline in comparison to walking training. Future cross-sectional and longitudinal research should also consider to account for a number of moderating factors (e.g., fitness level in cardiorespiratory, muscular and motor dimension, frequency, intensity and duration of relevant activities, whether activities are performed alone or with friends, etc.) and to evaluate additional neurobiological mechanisms (e.g., brain metabolism, muscle energetics, reduced oxidative stress, reduced psychological stress) through which Tai Chi may influence cognitive performance (Cheung et al., 2018; Gatts, 2008; Goon et al., 2009; Guo et al., 2014; Zhou et al., 2018).

## DATA AVAILABILITY STATEMENT

The datasets generated for this study are available on request to the corresponding author.

## ETHICS STATEMENT

Ethical approval for data collection and sharing was given by the Review Board of Soochow University. The participants provided their written informed consent to participate in this study.

## AUTHOR CONTRIBUTIONS

CLY and LYZ designed the study, CLY, ZHZ, JM and YLT collected and analyzed data, CLY, LYZ, YJZ, QY and JM wrote manuscript. All authors have read and agreed to the published version of the manuscript.

